# Incorporation of TRDV segments into TCR alpha chains

**DOI:** 10.1101/2023.02.26.530072

**Authors:** Michael Volkmar, Elham Fakhr, Stefan Zens, Rienk Offringa, Alice Bury, Jessica Gordon, Enes Huduti, Thomas Wölfel, Catherine Wölfel

**Affiliations:** Helmholtz Institute for Translational Oncology (HI-TRON) Mainz, 55131 Mainz, Germany; German Cancer Research Center (DKFZ), Department D200, 69120 Heidelberg, Germany; BioNtech, 55131 Mainz, Germany; Internal Medicine III, University Cancer Center (UCT), Research Center for Immunotherapy (FZI), University Medical Center (UMC) of the Johannes Gutenberg University Mainz and German Cancer Consortium (DKTK), Partner Site Frankfurt/Mainz, 55131 Mainz, Germany

## Abstract

In human tumor models we tried to identify and clone the TCR of tumor-reactive T cells enriched in mixed lymphocyte tumor-cell cultures (MLTC). In a particular MLTC, we identified a predominant TCR beta chain using the Beta Mark Vbeta Kit, but a corresponding alpha chain could not be amplified via RT-PCR using *TRAV*-specific forward primers. Therefore, we applied 5’RACE to obtain the TCR alpha chain sequences. The 5’RACE product revealed an alpha chain that encompassed 89bp of the *TRDV1* 5’UTR, followed by the *TRDV1* coding sequence joined in frame to *TRAJ24*. The ORF reaching from the *TRDV1* start codon to the *TRAC* segment was intact, suggesting a functional TCR. To analyze this MLTC population in greater depth we conducted 10X VDJ sequencing. CellRanger identified the beta chain known from the Beta Mark analysis, but no corresponding alpha chain in the filtered results. The corresponding TRDV-containing TCR alpha chain could, however, be detected in the “all_contig_annotations” files.

In a separate project, we performed TCR sequencing of tumor-infiltrating lymphocytes (TILs) in a murine tumor model. Also here, a predominant clonotype contained a TCR alpha chain joining *Trdv2-2* in frame to *Traj49*.

Transfection of both TCR cDNAs resulted in cell surface localization of TCR and CD3 as validated by FACS. Tumor recognition of the human, TRDV1-containing TCR could be demonstrated by IFNgamma ELISpot whereas the murine TCR did not recognize a tumor-derived cell line.

TRDV-containing alpha chains have been reported in the literature for two HLA I-restricted TCRs against HIV peptides (Ueno et al, Eur J Immunol, 2003). To determine whether such TDRV-containing TCRs are unique events or whether Vdelta segments are commonly incorporated into TCR alpha chains, we queried the NCBI Sequence Read Archive (SRA) for 10X VDJ data and analyzed 21 human and 23 murine datasets. We found that especially *TRDV1, Trdv1* and to some extent *Trdv2-2* are more commonly incorporated into TCR alpha chains than some *TRAV* genes, making the *TRDV* segments a relevant contribution to TCR alpha diversity.

For apparently solitary beta chains in 10X VDJ datasets, we suggest to scrutinize the “all_contig_annotations” files as these may contain an accompanying, TRDV-containing alpha chain.

## Introduction

The strength of adaptive immunity is its vast antigen receptor diversity that is generated by V(D)J recombination. Developing lymphoid cells express the recombinase proteins RAG1 and RAG2 that mediate the recombination of V and D or V and J genes. This recombination event creates the complementarity-determining region 3, CDR3, a crucial determinant of antibody or T-cell receptor specificity.

Usually, recombination between Vα, or *TRAV*, with Jα, or *TRAJ*, genes facilitates the generation of a specific TCRα chain in CD4^+^CD8^+^ αβ T cells whereas a joining of Vδ with Dδ and Jδ occurs in CD4^-^ CD8^-^ γδ T cells and produces the TCRδ chain. However, there have been identified “promiscuous” V genes that can be part of alpha as well as delta TCR chains; these are called TRAV/DV genes. In human, there are five known TRAV/DV genes (*TRAV14/DV4, TRAV23/DV6, TRAV29/DV5, TRAV36/DV7*, and *TRAV38-2/DV8*); in mouse there are nine such genes. In contrast, the *TRDV* genes are generally thought to recombine exclusively with Dδ and Jδ genes. Exceptions to this rule have been reported in nurse sharks [1] and in two TCRs isolated from a HIV patient. These Vδ-containing αβTCRs recognize HIV peptides presented by HLA class I molecules [2, 3].

In this work, we describe two TCRs, one of murine and one of human origin, that contain a Vδ segment in their TCRα chains. These TCRs represent the most abundant and the second-most abundant clonotypes in their respective TCR repertoires. Furthermore, to establish that *TRDV* genes are regularly incorporated into TCRα chains, we analyzed multiple publicly available TCR repertoire datasets and found Vδ-containing TCRα chains in the majority of them.

## Results

### 1st encounter of a TRDV-containing TCR alpha chain

For a donor PBMC-derived T cell culture that showed reactivity to a tumor cell line we tried to isolate the TCR that conferred the observed reactivity. We were able to identify the Vβ using the Beta Mark kit (Beckman Coulter, Brea, CA, USA) and to amplify the nucleotide sequence of the TCRβ chain using our standard amplification method [4]. However, the TCRα chain was refractory to this RT-PCR-based method using *TRAV*-specific forward primers. Therefore, we applied 5’RACE to amplify the TCRα chain sequence (*cf*. Material & Methods section and Table S1 for PCR primer sequences). The amplification was successful and Sanger sequencing of the 5’RACE product revealed a TCR chain that encompassed 89bp of *TRDV1* 5’UTR, followed by the *TRDV1* coding sequence which was joined to *TRAJ24*03* by a CALGDCITDSWGKFQF-encoding CDR3 sequence (Table 1). The open reading frame from the *TRDV1* start codon to the *TRAC* segment was intact, suggesting a functional T-cell receptor.

**Table 1.**
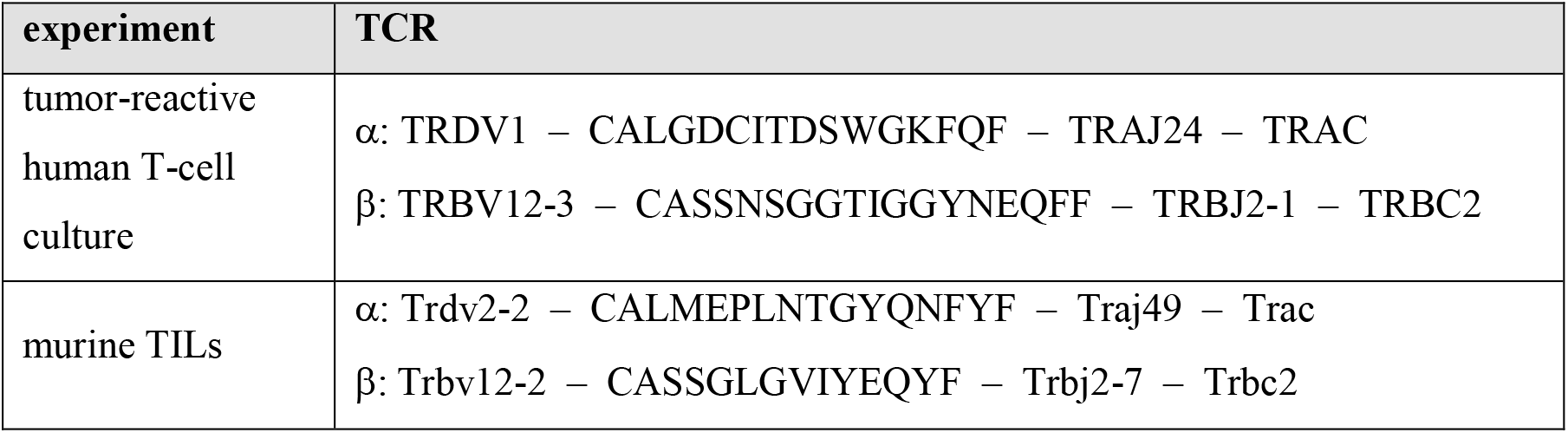
Composition of the TRDV-containing TCRs in this study.

To analyze this T cell culture in greater depth and identify further clonotypes we conducted 10X VDJ sequencing on T-cells from the same culture. The 10X CellRanger vdj pipeline identified the same TCRβ chain amplified before by RT-PCR for the second-most frequent clonotype of this T-cell culture (Table 1) but, seemingly, no TCRα chain. Manually inspecting the files generated by the 10X CellRanger pipeline we found no TCRα chain in the final, filtered results (`vloupe` file, `filtered_contigs.csv`, `web_summary.html` etc.). However, the entries for this clonotype in the “`all_contig_annotations.csv`” file showed *TRDV1* joined to *TRAJ24* as identified before by 5’RACE. Also the CDR3 sequence CALGDCITDSWGKFQF was identical to that derived from the 5’RACE product. The cell barcodes unambiguously linked this TCRα chain to the solitary TCRβ chain of the final, filtered 10X VDJ results. This TCR will be referred to as 29.ct2 subsequently.

### 2nd encounter, a TCR alpha chain containing Trdv2-2

In an independent experiment, we performed TCR sequencing of tumor-infiltrating lymphocytes, TILs, from a murine tumor model. The results indicated that the most frequent clonotype among the isolated TILs (clonotype 1) apparently contained no TCRα chain. As with the human T-cell culture described above, we manually inspected the “`all_contig_annotations.csv`” file generated by the 10X CellRanger vdj pipeline to search for a potential alpha chain that shared the cell barcodes of the apparently solitary beta chain. We found a TCRα chain that shared cell barcodes with the clonotype 1 beta chain and joined *Trdv2-2* to *Traj49* via a CDR3 region encoding CALMEPLNTGYQNFYF (Table 1). We refer to this TCR as m25800.ct1.

### TCR chains possess intact CDS

Both TCRα chains were categorized as non-productive by CellRanger (‘productive=false’ in results files), suggesting a disrupted open reading frame that does not encode an intact protein. However, since the 5’RACE results indicated a TCR chain with a functional coding sequence, we extracted the sequence assemblies from the respective “`all_contig_annotations.json`” files. We found that these sequences encode the V-segments (including 5’ untranslated region, UTR), CDR3 region, J-segment and 5’part of the C-segment all in-frame as an intact coding sequence. It can be inferred that the downstream part of the C-segment is also intact (for technical reasons, the downstream parts of the C-segments are not contained in 10X 5’libraries used for TCR sequencing, ref. [5]). Therefore, we concluded that the *TRDV*-containing TCRα chains contain complete, intact open reading frames.

### TCRs are expressed on the cell surface of T cells

The information in the “`all_contig_annotations`” files of both experiments enabled us to clone the TCRs for functional testing. To this end we used our previously established TCR cloning system [6, 7] that inserts V(D)J segments upstream of murine C segments that contain an additional cysteine residue facilitating a disulfide bridge between the alpha and beta C segments. The amino acid sequences of both recombinant TCRs are provided in Figure S1.

To test localization of the recombinant TCRs to the cell surface, we used JΔE10 cells, Jurkat E6.1-derived cells in which both TCR chains are disrupted by CRISPR/Cas9-introduced frameshift InDels in the respective C segment genes. The JΔE10 cells were transfected with *in vitro*-transcribed mRNA of the TCR constructs and analyzed by FACS. As depicted in Figure 1, JΔE10 cells transfected with TCR-encoding mRNAs show a strong signal for the murine TCR C segment whereas mock-transfected cells were not stained by this antibody. Figure 1 also shows that non-transfected JΔE10 cells are CD3-negative (left panel). Upon expression of the TCRs, the cells become CD3^+^, indicating a colocalization of TCR and CD3 at the cell surface.

**Figure 1.**
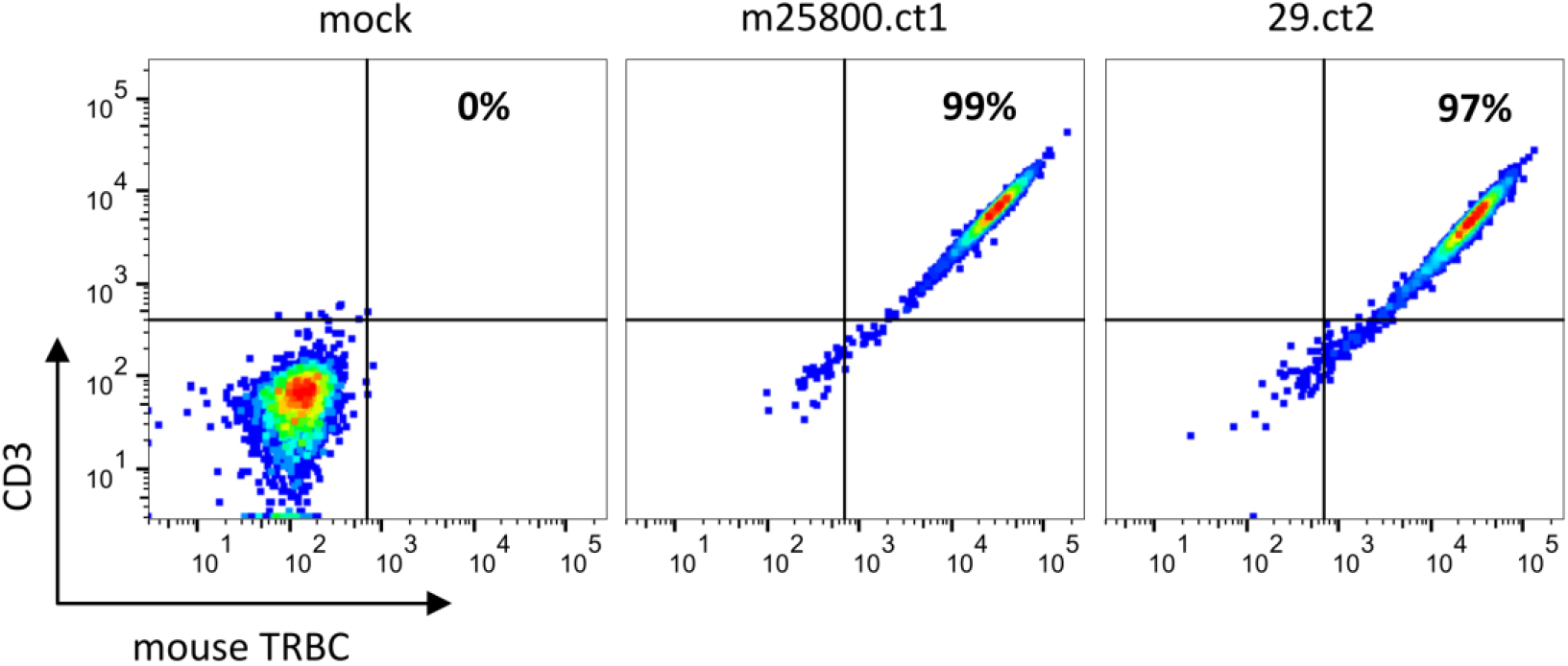
Flow-cytometric analysis depicting expression of recombinant TCRs on JΔE10 Jurkat cells 48h after transfection of *in vitro*-transcribed TCR mRNAs. The percentages in the upper right quadrant enumerate those JΔE10 cells that are double-positive for TCR and CD3. For each analysis 10,000 cells were acquired.

### T-cells transfected with TCR 29.ct2 recognize tumor cells

Despite the previously reported interaction of a Vδ-containing αβTCR with a MHC-peptide complex [2, 3], we could not assume *ab initio* that the TCRs we had identified were functional. Therefore, we introduced the 29.ct2 TCR into primary T-cells from a healthy blood donor after knocking out their endogenous TCR chains. Expression of the transgenic TCR was verified by FACS analysis. Subsequently, we used these T cells in an IFNγ ELISpot assay. Co-culture of TCR 29.ct2 transgenic T cells with the tumor cells resulted in very strong release of IFNγ, while no relevant IFNγ production was observed without tumor cells (Figure 2).

**Figure 2.**
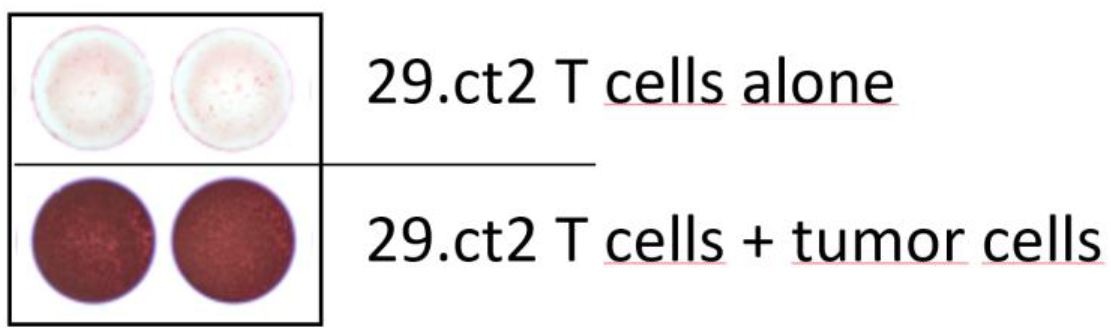
ELISpot assay of T-cells transduced with TCR 29.ct2. While the T-cells show very limited IFNg secretion if incubated without tumor cells (top panel), co-culture with tumor cells elicited a strong IFNγ signal (bottom panel).

The m25800.ct1 TCR was found to be non-reactive in a FACS-based assay measuring CD107a on transgenic T cells after co-culture with a cell line derived from the original mouse tumor.

Taken together this demonstrates that the TRDV1-containing TCR 29.ct2 confers binding of the T cells to the tumor cells and induces a very strong IFNγ response.

### TRDV-containing TCR alpha chains are common in human and mouse

Next, we aimed to determine whether the two TDRV-containing TCRs we discovered were rare events or whether Vδ segments are commonly incorporated into TCRα chains. To this end, we queried the NCBI Sequence Read Archive (SRA) for 10X VDJ data deposited in the form of Fastq files, downloaded and analyzed 43 randomly selected datasets, 21 of human and 22 of murine origin (Table S1).

To quantify the subset of Vδ-containing TCR clonotypes, we first extracted lines with the entry “TRDV” from the “`all_contig_annotations.csv`” file. Then, we quality-filtered the entries according to the following criteria:

- ≥2 UMIs (at least 2 Vδ-containing mRNAs in this cell have been reverse-transcribed and sequenced)
- high_confidence = true (the CellRanger software assigned a high confidence to this contig)
- full_length = true (the complete length of the targeted part of the open reading frame could be assembled from the sequencing reads)
- is_cell = true (the software deemed this contig to originate from an actual T-cell)
- The “productive” criterion was disregarded as the two TCRs we described above were deemed non-productive by the CellRanger versions used (≤7.0.1).

Then, based on the cell barcodes of the *TRDV*-containing TCRα chains, the other TCR chain(s) of the respective cells were extracted and the resulting dataset was split into two fractions: (i) cells that exclusively contained a Vδ-TCRα chain alongside a TCRβ chain and (ii) cells with more than one TCRα chain among which one contained a Vδ segment. For the first group, we manually copied the ‘raw_clonotype_id’ of the TCRβ chain to the TCRα chain with the same cell barcode because the Vδ-TCRα contigs were not assigned the clonotype ID by the CellRanger software. The second group, harboring *TRAV*-as well as *TRDV*-containing contigs was, again, manually filtered and all entries were deleted in which a “normal”, *TRAV*-containing, TCRα contig had an equal or higher UMI count. This left only those cells in which *TRDV*-TCRα contigs were more abundant than their *TRAV*-TCRα counterparts. The nonredundant *TRAV*-containing TCRα repertoire of each dataset was used as a control set for statistical and abundance determinations.

To obtain a comprehensively comparative picture on the *TRDV* usage in human TCRα chains, we merged the two curated subsets described above and the *TRAV*-TCRα repertoire of all 21 analyzed datasets, retaining the sample ID for identification and correct clonotype rank/size calculation. This yielded 71,232 TCRα clonotypes. We found *TRDV1* and *TRDV3* are incorporated into TCRα chains; we did not identify a *TRDV2*-containing TCRα chain that met our quality filter criteria. As shown in Figure 3A, *TRDV1* is not very frequently used in V-J recombination of the TCRα chain. However, it is more often used than four *TRAV* genes (*TRAV18, TRAV34, TRAV40* and *TRAV9-1*) while *TRAV18* and *TRAV9-1* are even more seldomly incorporated than *TRDV3*. When comparing clonotype sizes, for which we computed the mean rank of each V gene, *TRDV1* appears to be on par with the *TRAV* genes whereas *TRDV3* tends to be present in smaller clonotypes, only *TRAV9-1* being inferior in this respect (Fig. 3B). The minimal V gene rank shown in Fig. 3C indicates that *TRDV1* (top rank = 2, in our in-house dataset “line29”) and *TRDV3* (top rank =4, in public dataset SRR13113846) can become major clonotypes in TCR repertoires.

**Figure 3.**
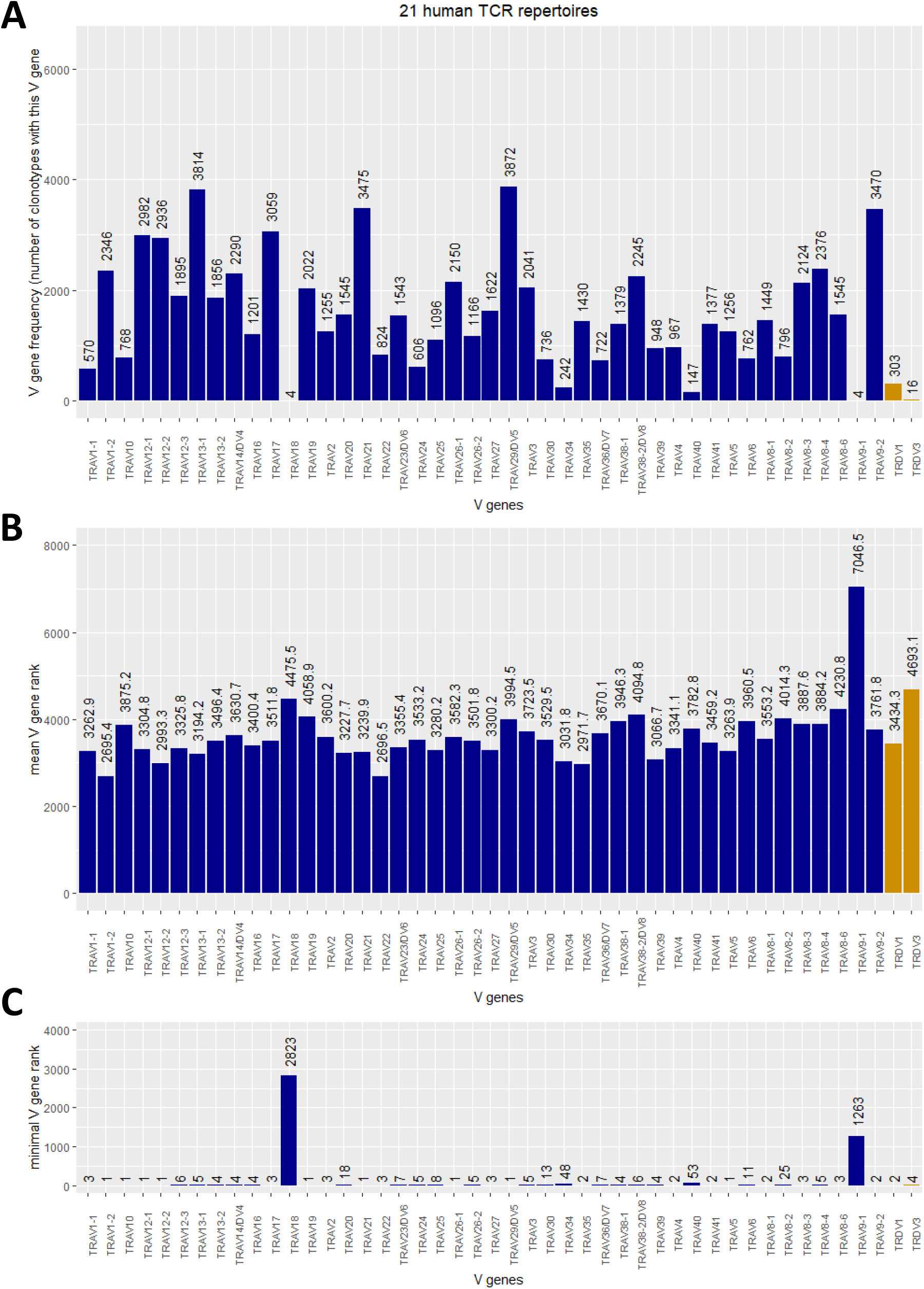
(A) Frequency plot of *TRAV* incl. *TRAV/DV* (dark blue) and *TRDV* (orange) genes in the TCRα chains of 21 analyzed human TCR repertoires. (B) mean rank of clonotypes carrying the respective V segment. (C) Minimal, or “top”, rank of a clonotype carrying the respective V gene among the 21 human TCR repertoires.

Merging the TCRα chains of the 22 mouse repertoires produced a table of 31,831 clonotypes. As shown in Figure 4, *Trdv1, Trdv2-1, Trdv2-2* and *Trdv5* are incorporated into murine TCRα chains. Although there are 75 *Trav* segments more abundant than *Trdv1*, the most often utilized *Trdv* segment, there are also 28 *Trav* genes that are incorporated less frequently (Fig. 4A). The frequency of utilization for *Trdv2-1* and *Trdv2-2* is in the same range as those of rarely incorporated *Trav* genes like *Trav12D2-2* and *Trav9D-3*, while *Trdv5* was found in only one clonotype. If, however, a *Trdv* segment is incorporated into TCRα chain, the corresponding clonotype can become big, shown as low minimal rank in Fig. 4C: m25800.ct1, the largest clonotype in our TIL sample, includes *Trdv2-2*; and two *Trdv1*-containing clonotypes are within the top10 most abundant clonotypes of their respective repertoires (top rank = 3 in dataset SRR18687603, top rank = 5 in SRR18687609).

**Figure 4.**
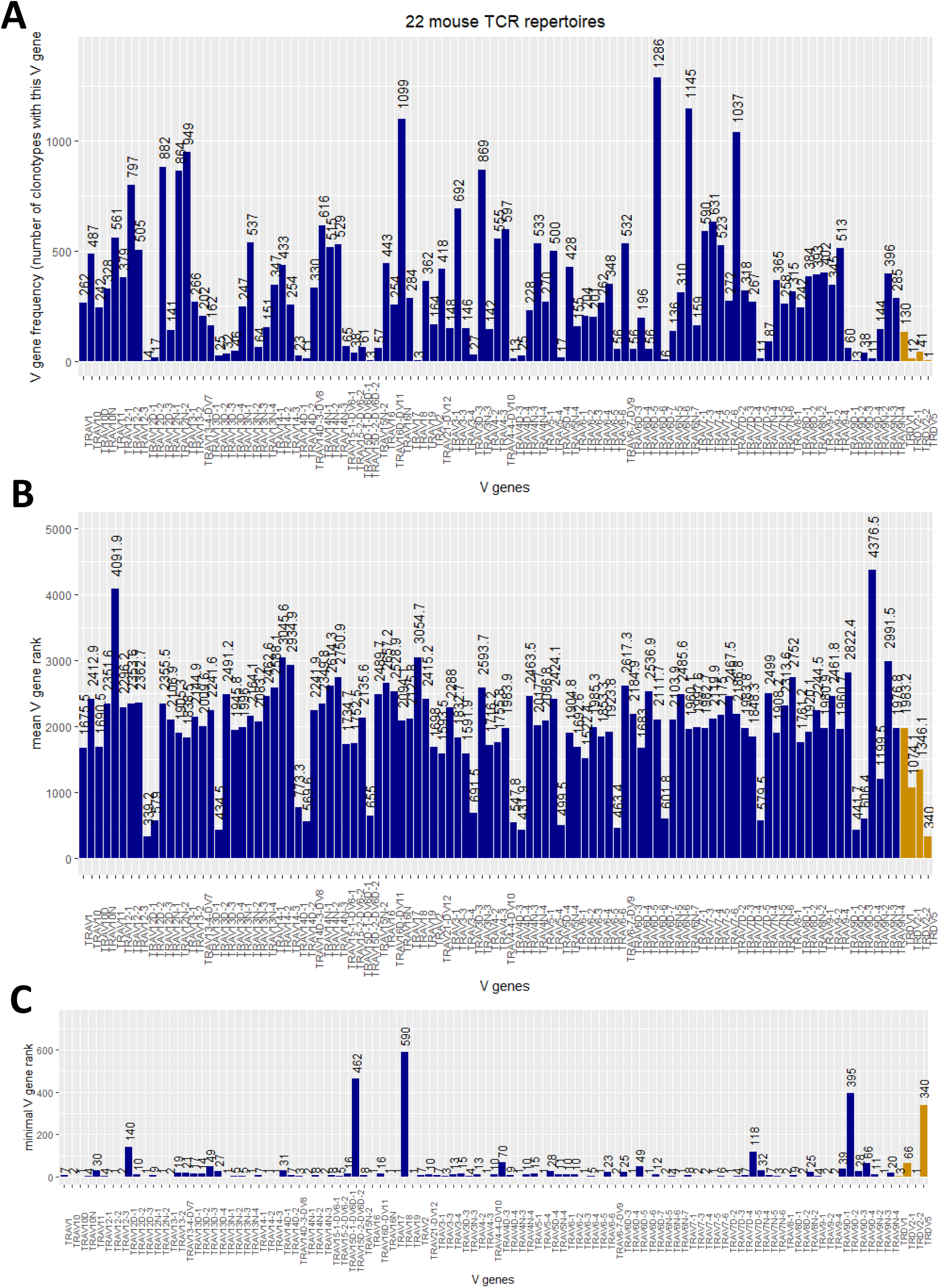
(A) Frequency plot of *Trav* and *Trdv* genes in the TCRα chains of 22 analyzed murine TCR repertoires. (B) mean rank of clonotypes carrying the respective V segment. (C) Minimal, or “top”, rank of a clonotype carrying the respective V gene among the 22 mouse TCR repertoires.

*Trdv4* occurred in only one cell barcode in dataset SRR21989196. But in the same cell, a more abundant *Trav13D1*-containing TCRα chain was present (5 and 9 UMIs, respectively), so this chain did not meet our criteria. Nonetheless, there might be TCR repertoires in which *Trdv4* and, although we did not identify it in our screen, *Trdv3* are productively incorporated into TCRα chains.

Taken together our combined analyses of multiple human and mouse TCR repertoires demonstrated that *TRDV* gene segments are incorporated into TCRα chains at levels exceeding those of some *TRAV* genes. The sizes of the resulting clonotypes are on average comparable with those of *TRAV*-containing TCRs and can become large, ranking in the top10, or even dominate the repertoire as in our samples 29 and m25800.

### J gene usage in TRDV-TCRα and TRAV-TCRα chains is similar

We were wondering whether there was a bias in the *TRAJ* utilization for *TRAV*-TCRα and *TRDV*-TCRα recombination events. As the number of individual *TRDV* gene-containing TCRα chains is too small for a meaningful distribution analysis, we analyzed *TRAJ* frequencies of all *TRDV*-containing *versus* all *TRAV*-containing TCRα chains but found no statistically significant differences among human and mouse TCRs (X^2^ test p-value 0.25 in both cases). The *TRAJ* frequency plots are shown in Fig. S2. Interestingly, among the murine TCRs there appears to be a tendency to utilize more downstream located *TRAJ* genes, independently of whether the V segment is a Vα or a Vδ gene (Fig. S2 bottom panel) whereas in the utilization frequency of human *TRAJ* genes this trend seems to be overlaid several dips in frequency *e*.*g*. between *TRAJ24*-*TRAJ26, TRAJ32*-*TRAJ36*, and *TRAJ45*-*TRAJ48*.

## Discussion and Outlook

In this work, we identified two highly enriched TCR clonotypes that contain *TRDV* segments in their TCR alpha chains. Upon transfection into T cells, the TCRs are translocated to the cell surface and the cells also become CD3-positive, suggesting the establishment of a TCR-CD3 complex on the cell membrane. While we could verify tumor reactivity of the 29.ct2 TCR by ELISpot, the m25800.ct1 TCR was not reactive against an autologous tumor cell line. Explanations for the non-reactivity could be that the mutation evoking the expansion of the m25800.ct1 T cells in the tumor is either not contained or epigenetically silenced in the tumor-derived cell line or that this was a non-reactive bystander clonotype. Despite the two TCRs being highly expanded in their samples, representing the top1 and top2 TCR respectively, these could have been unique cases that, while interesting in themselves, bear no practical relevance for TCR repertoire analysis. Drawing from over 40 public, high-throughput single cell TCR sequencing datasets, we showed that *TRDV*-containing TCRα chains are present in the vast majority of repertoires. Actually, the datasets lacking this kind of alpha chain tend to be smaller and more oligoclonal. This suggests that utilization of *TRDV* genes in V(D)J recombination to generate TCRα chains is not an exception but rather common. Our analyses demonstrated that *TRDV* genes are not as frequently used as many *TRAV* genes, representing roughly 0.45% among human and 0.6% among murine TCRα clonotypes. However, especially *TRDV1, Trdv1* and to some extent *Trdv2-2* are more commonly incorporated into TCRα chains than some *TRAV* genes, making the *TRDV* segments a relevant contribution to TCRα diversity. Moreover, we observed *TRDV*-TCRα chains among the top10 clonotypes not only in our datasets but also public ones.

The recombination of *TRDV*, or *TRDD*, to *TRAJ* segments does not violate the 12/23 rule [8] and has been proven experimentally to some extent. However, utilization of V genes in TCRα recombination is thought to be governed at least in part by features of their promoters and chromatin accessibility [9, 10]. While we found a tendency towards utilization of more downstream located *TRAJ* genes, this was present in both, *TRAV*-TCRα as well as *TRDV*-TCRα chains and we observed no apparent difference between the two types of alpha chains.

Our results give rise to new questions: Can, for example, gamma-delta T-cells harbor *TRAV* segments in their TCRδ chains? Another interesting aspect is the apparent co-occurrence of “normal” *TRAV*-TCRα chains in the same cells that carry *TRDV*-TCRα chains where the UMI count suggests that both are expressed. Do those T-cells display two distinct types of TCR on their surface and are both functional? We hope that the work presented in this manuscript will establish the incorporation of *TRDV* genes into functional human and murine TCRs as a verifiable fact thus facilitating work on such questions as well as generally broadening the field of TCR repertoire analysis.

It may happen to our fellow colleagues working with TCR repertoire data obtained from 10X VDJ sequencing that they encounter TCR clonotypes that seem to contain only a TCR beta chain with no accompanying alpha chain. For those cases we humbly suggest to turn to the unfiltered “`all_contig_annotations`” files. More often than not, these files may harbor a TRDV-containing

TCR alpha chain sharing the cell barcode(s) of the apparently solitary beta chain. This TRDV-TCRα chain may, together with the beta chain, constitute a functional T cell receptor.

## Material & Methods

### TCR 5’RACE

Total RNA was isolated from a tumor-reactive T cell culture using a Qiagen RNeasy Micro kit (Qiagen GmbH, Hilden, Germany). 1μg total RNA was used for reverse transcription with the NEB Template Switching Reverse Transcriptase Enzyme Mix (New England Biolabs, Ipswich, MA, USA) according to the recommendations of the manufacturer. We used an oligonucleotide specific for the human TCR alpha chain C segment (hsTRAC-RT, see Table S2) to prime the reverse transcriptase, and a template-switching oligonucleotide (TSO) to create a generic primer binding site at the 5’ends of synthesized cDNAs. The RT product was subjected to 2 rounds of PCR amplification. All oligonucleotides used were obtained from Merck Sigma-Aldrich GmbH (Darmstadt, Germany). The product of the nested PCR was cloned into pJET (Thermo-Fisher, Waltham, MA, USA) and four isolated plasmid clones were subjected to Sanger Sequencing (Eurofins Scientific SE, Luxembourg, Luxembourg). Analysis of the resulting sequences was done using IMGT/V-QUEST [11].

### 10X TCR sequencing

Number and viability of isolated T-cells was assessed either by manual counting with Trypan Blue or using a Casy TT cell counter (OMNI Life Science GmbH & Co KG, Bremen, Germany). VDJ sequencing libraries were established according to the manufacturer’s protocols (10X Genomics, Pleasanton, CA, USA). Paired-end sequencing was performed either in-house (HI-TRON Mainz, Mainz, Germany) on an Illumina MiSeq or by the DKFZ Genomics Core Facility (Heidelberg, Germany) on a NovaSeq6000, aiming for ≥5,000 reads/cell. Fastq files were analyzed using the CellRanger vdj pipeline as recommended by the manufacturer (*cf*. Data Analysis).

### Cloning of TCRs and in vitro Transcription

We ordered the variable parts (V-CDR3-J) of the TCR chains as synthetic dsDNA oligonucleotides from Twist Bioscience (San Francisco, CA, USA) and cloned them in-frame with murine alpha/beta C-segments that are stabilized by an additional disulfide bridge [12, 13] using Golden Gate Assembly as described earlier [7] into a pcDNA3.1(+) backbone; the pcDNA3.1 plasmid was purchased from Thermo-Fisher (Waltham, MA, USA). After sequence verification via Sanger sequencing, 1μg of NotI-linearized plasmid was subjected to *in vitro* transcription using the HiScribe T7 ARCA mRNA Kit (with tailing) (New England Biolabs, Ipswich, MA, USA). For use in ELISpot, the 29.ct2 TCR was cloned as described in [6].

### Electroporation of modified Jurkat cells

Jurkat E6-1 were obtained from ATCC (Manassas, VA, USA) and grown in RPMI-1640 with 10% FBS and 1% Non-Essential Amino Acids solution (Thermo-Fisher). The endogenous TCR chains of the Jurkat cells were knocked out using Crispr/Cas9 targeting the first exon of the TCR C segments.

For electroporation of TCR constructs, 1×10^6^ cells were suspended in 20μl Opti-MEM medium (Thermo-Fisher) and 2.5μg of IVT-RNA were added. For the mock control, the cells were only suspended in Opti-MEM. Cells and RNA were mixed by gentle pipetting, transferred into a Nucleocuvette and immediately electroporated using the CL120 program in the 4D-Nucleofector X Unit (Lonza, Walkersville, MD, USA). After electroporation, 80μl pre-warmed complete RPMI medium (37°C) were added and the cells were let to rest for 10min. Then, the cell suspension was transferred to 1ml pre-warmed complete RPMI medium (48-well plate) and incubated at 37°C, 5% CO_2_.

### FACS analysis

48 hours after electroporation, cells were analyzed for TCR expression. The cells were first washed using pre-chilled Cell Staining Buffer (Biolegend, San Diego, CA, USA) supplemented with 2mM EDTA. Then, the cells were stained with LIVE/DEAD Fixable Green Dead Cell Stain Kit (Thermo-Fisher) in a 1:1000 dilution for 30min on ice. The samples were washed with Cell Staining Buffer and stained with anti-mouse TCR β chain-Alexa Fluor^®^ 647 (Biolegend) at a concentration of 0.02μg/μl, for 30min on ice. Then, the cells were washed and fixed with Fixation Buffer (Biolegend) mixed with an equal volume of Cell Staining Buffer and incubated for 30min at room temperature. Finally, cells were passed through a 30μm filter and analyzed using a BD FACSAria™ Fusion instrument. Unless indicated otherwise, 10,000 cells were analyzed per condition.

### ELISpot assay

IFNγ-ELISpot assays were performed as described previously [6, 14]. Briefly, tumor cells were seeded (50,000 cells/well) into ELISpot plates (Millipore, Burlington, MA, USA). TCR-transfected T-cells (10,000 T-cells/well) were incubated with or without tumor cells for 20-24h. Subsequently, the plates were developed and imaged using an ImmunoSpot analyzer device (Cellular Technology Limited, Cleveland, OH, USA).

### Data analysis and visualization

After testing several versions of the CellRanger software, raw 10X VDJ sequencing datasets were analyzed with CellRanger v7.0.1 utilizing the GRCh38-7.1.0 human or the GRCm38-7.0.0 murine reference, respectively. Further analyses and visualizations were performed with R 4.2.0 [15] using the following packages: ggplot2 [16], dplyr [17] and knitr [18, 19]. FACS scatterplots were exported as SVG files and modified with Inkscape (v1.0) to improve clarity.

## Author contributions

M.V. conceptualized the work. C.W., E.F., S.Z., A.B., J.G., E.H. and M.V. performed experiments and collected data. C.W., E.F. and M.V. analyzed and interpreted data. M.V. drafted the article. C.W., T.W. and E.F. critically reviewed the manuscript.

## Acknowledgements

We thank Ed Green (DKFZ) for valuable suggestions to improve the quality of this manuscript. We thank Martina Fatho for technical assistance. We thank the NGS Core Facility, German Cancer Research Center (DKFZ), for providing excellent NextGen Sequencing services.

## Supplementary Data

Table S1: TCR datasets used in this study

Table S2: PCR primers

Figure S1: Amino acid sequences of the TCR expression constructs

Figure S2: TRAJ frequency plots

