## Supplementary Tables and Figures for "Incorporation of TRDV segments into TCR alpha chains"

**Table S1.** 10X TCR scRNA datasets used in this study.

| human | murine |
|---|---|
| SRR13113571 | SRR10723081 |
| SRR13113572 | SRR10723082 |
| SRR13113573 | SRR10723083 |
| SRR13113574 | SRR10723084 |
| SRR13113575 | SRR14911004 |
| SRR13113576 | SRR18687603 |
| SRR13113577 | SRR18687605 |
| SRR13113578 | SRR18687606 |
| SRR13113579 | SRR18687607 |
| SRR13113580 | SRR18687608 |
| SRR13113581 | SRR18687609 |
| SRR13113582 | SRR18687610 |
| SRR13113846 | SRR18687611 |
| SRR15597465 | SRR18687613 |
| SRR18810015 | SRR18687614 |
| SRR18810016 | SRR18687616 |
| SRR18810018 | SRR18687617 |
| SRR20751884 | SRR18687618 |
| SRR20751886 | SRR18687619 |
| SRR20751888 | SRR21738977 |
| line29 | SRR21989196 |
| | m25800 |

**Table S2.** Primers used in this study.

| Primer name | Primer sequence | Purpose |
|---|---|---|
| TSO | AAGCAGTGGTATCAACGCAGAGTACATrGrGrG * | template-switching oligo during reverse transcription |
| TSO_preampPCRpr | AAGCAGTGGTATCAACGCAGAGTA | adapter-sepcific primer, 1st round of 5'-RACE-PCR |
| TSO_nestedPCRpr | GCAGTGGTATCAACGCAGAGTACAT | adapter-sepcific primer, nested RACE-PCR |
| hsTRAC-RT | TTCGGAACCCAATCACTG | reverse transcription of TCR alpha chain |
| hsTRAC-R1 | GCTTGACATCACAGGAACTTTCTGG | 1st round of 5'-RACE-PCR |
| hsTRAC-R2 | TGCTCTTGAAGTCCATAGACCTCATG | nested RACE-PCR |

* ‘r’ denotes reibonucleotides

TRDV1-containing TCR 29.ct2

MDSWTFCCVSLCILVAKHTDAGVIQSPRHEVTEMGQEVTLRCKPISGHNSLFWYRQTMMRGLELLIYFNNNVPIDDSG
MPEDRFSAKMPNASFSTLKIQPSEPRDSAVYFCASSNSGGTIGGYNEQFFGPGTRLTVLEDLRNVTPPKVSLFEPSKA
EIANKQKATLVCLARGFFPDHVELSWWVNGKEVHSGVCTDPQAYKESNYSYCLSSRLRVSATFWHNPRNHFRCQVQFH
GLSEEDKWPEGSPKPVTQNISAEAWGRADCGITSASYQQGVLSATILYEILLGKATLYAVLVSTLVVMAMVKRKNSGS
GATNFSLLKQAGDVEENPGPGMLFSSLLCVFVAFSYSGSSVAQKVTQAQSSVSMPVRKAVTLNCLYETSWWSYYIFWY
KQLPSKEMIFLIRQGSDEQNAKSGRYSVNFKKAAKSVALTISALQLEDSAKYFCALGDCITDSWGKFQFGAGTQVVVT
PDIQNPEPAVYQLKDPRSQDSTLCLFTDFDSQINVPKTMESGTFITDKCVLDMKAMDSKSNGAIAWSNQTSFTCQDIF
KETNATYPSSDVPCDATLTEKSFETDMNLNFQNLSVMGLRILLLKVAGFNLLMTLRLWSS*

Trdv2-2-containing TCR m25800.ct1

MSNTAFPDPAWNTTLLSWVALFLLGTSSANSGVVQSPRYIIKGKGERSILKCIPISGHLSVAWYQQTQGQELKFFIQH
YDKMERDKGNLPSRFSVQQFDDYHSEMNMSALELEDSAVYFCASSGLGVIYEQYFGPGTRLTVLEDLRNVTPPKVSLF
EPSKAEIANKQKATLVCLARGFFPDHVELSWWVNGKEVHSGVCTDPQAYKESNYSYCLSSRLRVSATFWHNPRNHFRC
QVQFHGLSEEDKWPEGSPKPVTQNISAEAWGRADCGITSASYQQGVLSATILYEILLGKATLYAVLVSTLVVMAMVKR
KNSGSGATNFSLLKQAGDVEENPGPGMVRPFFLWVLFLSTSLEASMAQTVSQPQKKKSVQVAESATLDCTYDTSDTNY
LLFWYKQQGGQVTLVILQEAYKQYNATLNRFSVNFQKAAKSFSLEISDSQLGDAATYFCALMEPLNTGYQNFYFGKGT
SLTVIPNIQNPEPAVYQLKDPRSQDSTLCLFTDFDSQINVPKTMESGTFITDKCVLDMKAMDSKSNGAIAWSNQTSFT
CQDIFKETNATYPSSDVPCDATLTEKSFETDMNLNFQNLSVMGLRILLLKVAGFNLLMTLRLWSS*

X VDJ beta chain, CDR3 sequence is underlined
X beta C segment
X P2A „self-cleaving“ sequence
X VDJ alpha chain, CDR3 sequence is underlined
X alpha C segment

**Figure S1.** Amino acid sequences of analyzed TCRs.

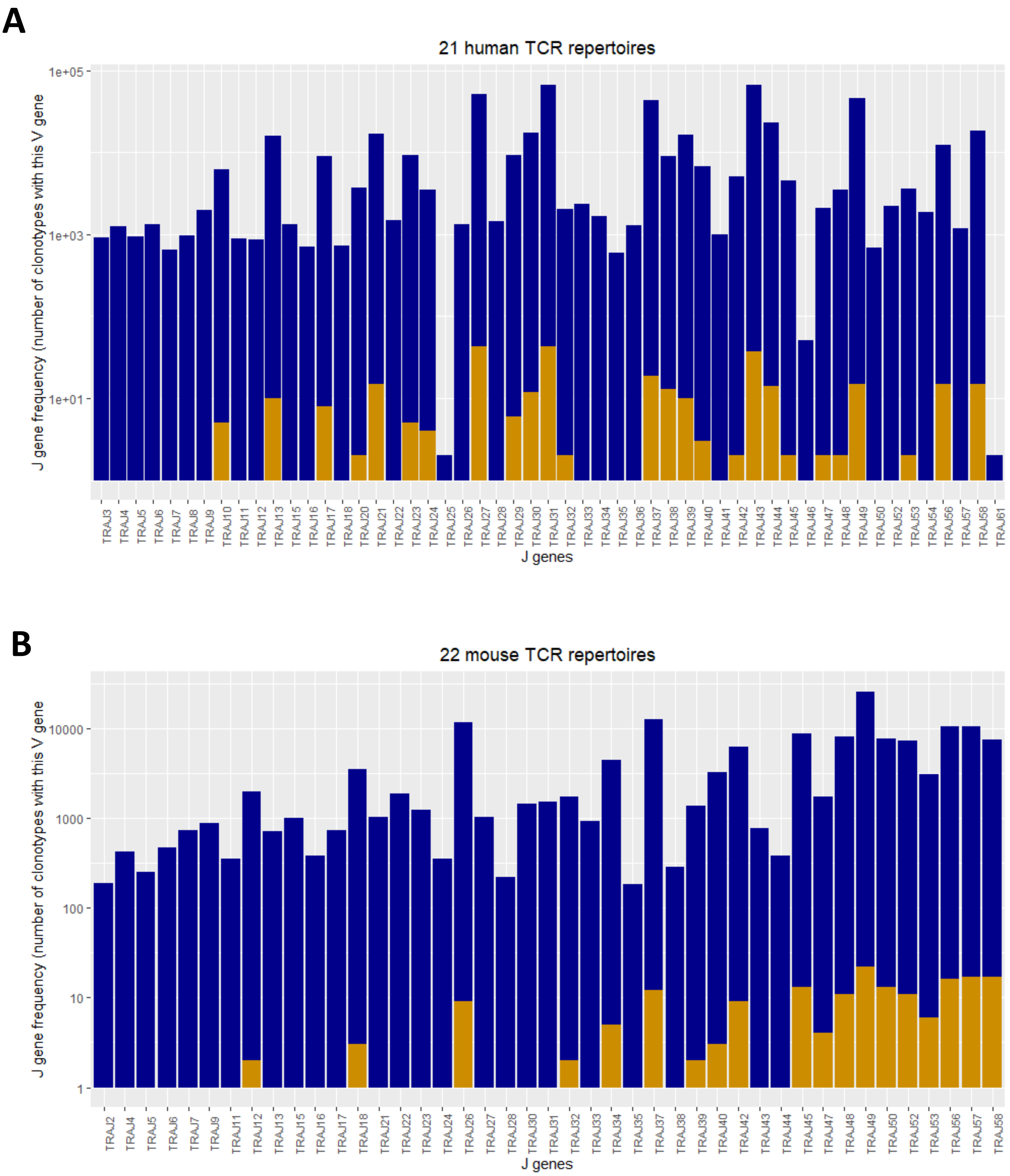


**Figure S2.** *TRAJ* frequency distribution among *TRAV*-containing (dark blue) and *TRDV*-containing (orange) TCRα chains. Accumulated data of all human TCRα repertoires is shown in (A), the murine repertoires are shown in (B).
